# The *ThCre:mTmG* mouse has sparse expression in the sympathetic nervous system

**DOI:** 10.1101/2023.02.01.526373

**Authors:** Tina Tian, Patricia J Ward

## Abstract

The global double-fluorescent cre-dependent reporter mouse *mT/mG* has been a valuable tool for differential identification of tissues and cells. This line was previously shown to have EGFP expression in a cre-dependent fashion in single neurons. Here, we demonstrate very sparse labeling in a *ThCre:mT/mG* cross. This is evident in sweat glands, thoracic spinal cord, sciatic nerve, and lumbar sympathetic ganglia. Therefore, this model may be suitable for imaging or electrophysiology studies of postganglionic sympathetic neuron cell bodies but ineffective for visualizing peripheral axons and tissue targets of the sympathetic nervous system.

## DESCRIPTION

Methods for differentially identifying subtypes of neurons is important in the neurosciences. For example, in the field of neural regeneration, understanding how novel therapeutics affect different populations of neurons would be incredibly useful for preferentially targeting the growth or inhibition of specific neurons after nerve injury (Baron 2000, Bennett 1991, Brennan et al 2021, Chung et al 1996, Haase & Chung 2002, Walters 2018, Ward et al 2018). The *mt/mg* mouse is a double-fluorescent cre-dependent reporter that expresses the membrane-targeted tandem dimer Tomato (tdTomato, mT) prior to cre-mediated excision and membrane-targeted enhanced green fluorescent protein (EGFP, mG) after excision (Muzumdar et al 2007). This mouse has been shown to have excellent conversion to EGFP expression with global cre drivers that are both constitutively active and tamoxifen-inducible. In the original article, the following tissues were evaluated: brain, retina, heart, lung, rib, liver, spleen, kidney, bowel, ovary, cerebellum, and olfactory bulb (Muzumdar et al 2007). Although fluorescent labeling of neuronal somas and processes was robust in culture of primary neurons from embryonic cortical caps and olfactory bulb sensory neurons in P21 mice, adult neuronal tissue notably had a marked lack of distinct cellular labeling.

The purpose of creating this *ThCre:mTmG* cross was to create a nerve injury model in which regenerating sympathetic axon profiles could be visualized in whole nerves with mG labeling in contrast to motor and sensory axon profiles, which would be expected to remain labeled with mT, without reliance on immunohistochemistry. However, in the bilateral L2-L5 lumbar sympathetic ganglia of *ThCre:mT/mG*, only 31.75 ± 8.77 (mean ± standard deviation) postganglionic sympathetic neurons exhibit mG labeling **(Fig. 1A-G)**. Given that mouse nerve composition is similar to that of the rat, the typical number of sympathetic axons in the sciatic nerve is approximately 6,210, or 12,420 bilaterally (Schmalbruch 1986). Therefore, labeling in the L2-L5 lumbar sympathetic ganglia is estimated to be 0.255%.

**Figure 1:**
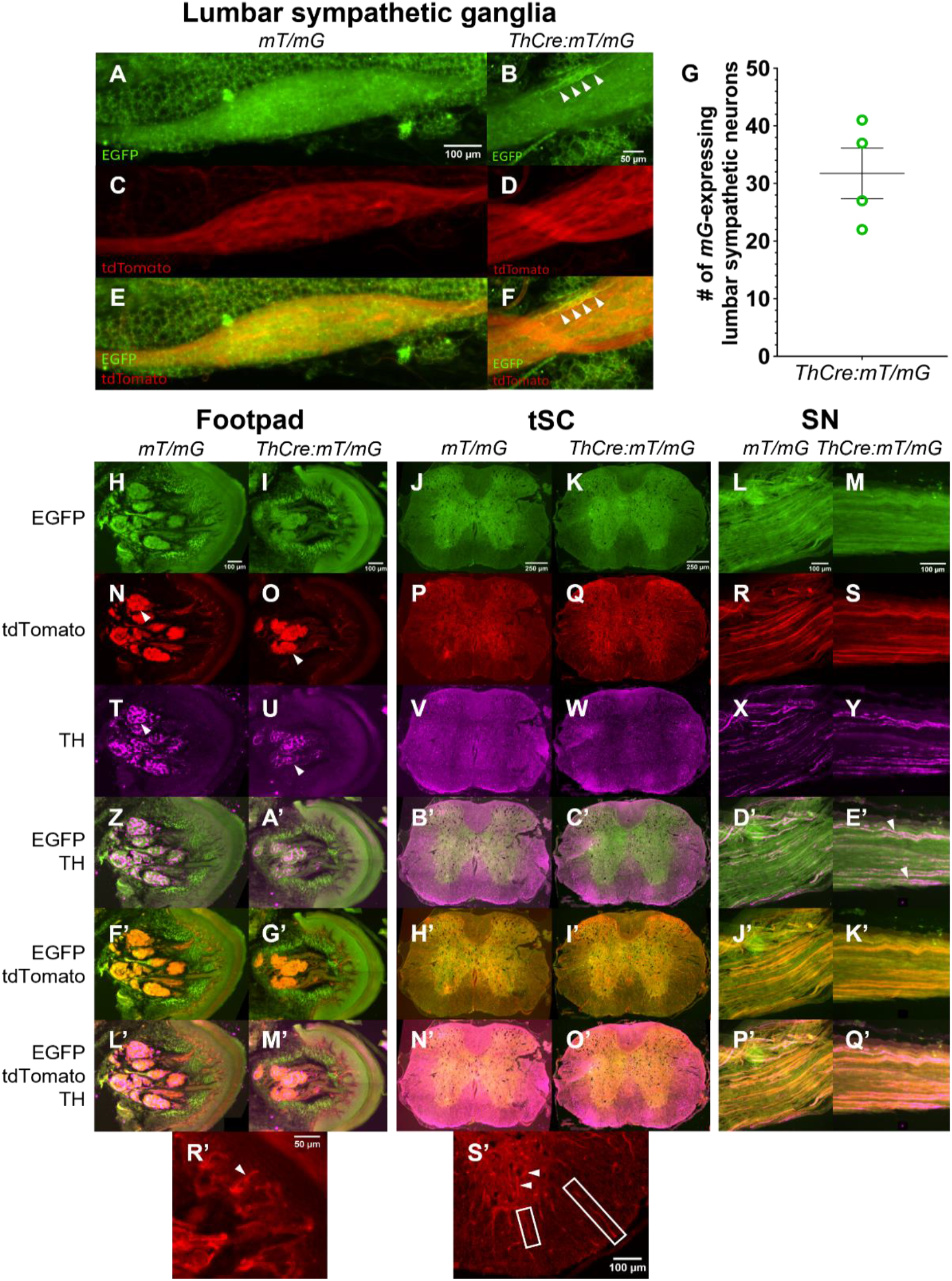
Expression of *ThCre*-induced EGFP in *ThCre:mT/mG* mice. (A-F) Expression of EGFP and tdTomato in the lumbar sympathetic ganglia of *mT/mG* mice (left) and *ThCre:mT/mG* mice (right). (B, F) EGFP-labeled sympathetic neurons (arrows). (G) Number of EGFP-expressing neurons in the bilateral L2-L5 lumbar sympathetic ganglia of *ThCre:mT/mG* mice, mean ± SEM. n = 4. (H-M) EGFP expression in the footpad, thoracic spinal cord (tSC), and sciatic nerves (SN) of *mT/mG* mice (left columns) and *ThCre:mT/mG* mice (right columns). (N-S) tdTomato expression. (T-Y) Tyrosine hydroxylase (TH) expression. (N, O, T, U) Sweat glands with TH innervation (arrows). (Z-E’) EGFP and TH merged. (E’) No overlap of TH+ axons and EGFP-labeled axons (arrows). (F’-K’) EGFP and tdTomato merged. (L’-Q’) EGFP, tdTomato, and TH merged. (R’) Magnified image of tdTomato-labeled dermal papillae structures seen in the footpad. Tubular structure in dermal papilla (arrow). (S’) Magnified image of tdTomato-labeled structures in the white and gray matters of the thoracic spinal cord with tubular structures within the gray matter (arrows) and axon-like structures in the white matter (boxes).

In the sweat glands of the footpads, thoracic spinal cord, and sciatic nerve, tyrosine hydroxylase-positive (TH+) neuronal labeling with mG/EGFP is negligible **(Fig. 1Z-E’)**. There is no difference in the green channels between *mT/mG* control mice versus *ThCre:mT/mG* mice **(Fig. 1H-M)**. Despite undetectable norepinephrine production within the axons that innervate sweat glands due to their cholinergic nature, mouse nerve fibers that innervate sweat glands still exhibit robust TH immunoreactivity visible via immunohistochemistry and are expected to express EGFP **(Fig. 1H-I, Fig. 1T-U)** (Landis et al 1988).

Additionally, mT/tdTomato labeling in the nervous system structures that have been analyzed to-date appears to be sparse at best. In footpad sections, sensory end organs are not visible in the dermal papillae – Instead, the structures that extend into the dermal papillae appear to form cylindrical structures, more akin to vasculature or sweat gland ducts **(Fig. 1R’)**. It is notable that the TH+ fibers present in the sweat glands are also not visible with mT labeling in *mT/mG* control mice **(Fig. 1N)**. mT labeling of lumbar sympathetic neurons also appears negligible in the control *mT/mG* mouse **(Fig. 1C)** as the fluorescent reporter is expected to fill the cell bodies of neuronal tissues.

In the spinal cords of *mT/mG* mice, axon-like structures with visible tdTomato-expression are more apparent in the white matter while in the gray matter, structures that resemble vasculature are predominant with no visible neuronal processes throughout the gray matter **(Fig. 1S’)** and no visible motoneuron cell bodies in the ventral horn **(Fig. 1P, 1Q)**. Finally, in the sciatic nerve, although axon-like structures are clearly visible with TdTomato-labeling **(Fig. 1R, 1S)**, the labeling is not ubiquitous. In comparison to neuronal fluorescent reporter mice such as the Thy-1 YFP-16 mouse, which exhibits robust fluorescence in motoneurons and sensory neurons (Feng et al 2000), the labeling of axons in the sciatic nerve with mT is sparse.

These results suggest that the *ThCre:mT/mG* mouse an ineffective model for visualizing structures of the sympathetic nervous system. Additionally, the *mT/mG* mouse itself exhibits sparse mT labeling in several other nervous system tissues. This finding can be indicative of high background fluorescence of other tissues that makes visualization of fine neuronal processes in tissue, compared to the original robust labeling in culture (Muzumdar et al 2007), extremely difficult. Although *mT/mG* mice in these assays show strong expression of mT in tissues such as the sweat glands themselves, vasculature-like structures, and epidermal and dermal skin structures, mT expression in neuronal structures is incomplete. Incomplete or sparse labeling may be advantageous for experiments involving high-resolution imaging, so this model can be very valuable in that respect. Additionally, non-neuronal tissue expression of mT is exceptional, which may make this mouse an optimal model for more diverse studies.

## METHODS

### Animals

Adult *mT/mG* mice were obtained from The Jackson Laboratory (strain #007576) (Muzumdar et al 2007), and *Th(Th-cre)Fl12Gsat/Mmucd* mice, referred to as *ThCre*, were obtained from the Mutant Mouse Resource and Research Centers (strain #017262-UCD) (Gong et al 2007, Gong et al 2003). Mutant *ThCre*:*mT/mG* mice were generated from crossing *ThCre* transgenic mice and *mT/mG* mice. *Th* (tyrosine hydroxylase) codes for the rate-limiting enzyme of catecholamine synthesis present in sympathetic neurons (Molinoff & Axelrod 1971).

All mice were 8-12 weeks of age and weighed between 15-25g. Groups contained equal males and females. A 25-week female *mT/mG* mouse, one of the founding breeders, from The Jackson Laboratory was also evaluated. All experiments were approved by the Institutional Animal Care and Use Committee of Emory University

### Immunohistochemistry

Mice were deeply anesthetized via intraperitoneal injection of 150 mg/kg 10 mg/ml Euthasol (0.75ml Euthasol [Virbac AH Inc, Fort Worth, TX, ANADA 200-071, NDC 051311-050-01] in 19.75ml 0.9% bacteriostatic sodium chloride [Hospira, Inc., Lake Forest, IL, NDC 0409-1966-02]) with a 25-gauge 1 mL syringe (BD, 309626). Mice were exsanguinated with 0.9% NaCl (Fisher Scientific, S2711) then perfused with 4% paraformaldehyde (Sigma-Aldrich, P6148) in 0.01M (1X) phosphate buffered saline (PBS, Sigma-Aldrich, P4417). Right plantar skin, right sciatic nerve, T10-T12 spinal cord, and the bilateral L2-L5 sympathetic ganglia were harvested and placed in 20% sucrose for cryoprotection and allowed to sink at 4°C prior to cryosectioning. Plantar skin was sectioned transversely at 30 μm to see footpad architecture, sciatic nerve longitudinally at 20 μm, and spinal cord transversely at 20 μm. Sympathetic ganglia were whole-mounted to a slide with Fluoro-Gel with Tris Buffer (Electron Microscopy Sciences, 17985-10).

Slides were blocked with 10% normal goat serum (VWR International, GTX73206) in 1X PBS for 1 hour at room temperature (RT) in a humidity chamber. Primary antibody rabbit anti-tyrosine hydroxylase (Abcam Ab112) was diluted at 1:750 in the blocking buffer and allowed to incubate overnight at RT. Slides were then washed with 1X PBS for 3 10-minute washes before applying the secondary antibody goat anti-rabbit Alexa 647 (Invitrogen A21245), which was allowed to incubate at RT for 2 hours. After washing with 1X PBS for 3 10-minute washes, slides were dried prior to mounting with Fluoro-Gel.

### Imaging

Slides were imaged on a Nikon Ti-E fluorescent microscope. All samples were imaged with a 20x objective. At least 3 sections of footpads, 3 sections of thoracic spinal cord, 2 sections of sciatic nerve, and the whole L2-L5 bilateral sympathetic ganglia were evaluated per animal. Exposure times as follows: TRITC 50 ms, FITC 400 ms, Cy5 200 ms. Images stitched with “ND Acquisition Large Images” function as needed.

### Analysis

Images were analyzed in Fiji, looking for overlap of TH+ fibers and cre-dependent mG expression. The number of mG-positive cells in the lumbar sympathetic ganglia in *ThCre:mTmG* mice was also counted.

